# Validation of non-invasive body-surface gastric mapping for detecting electrophysiological biomarkers by simultaneous high-resolution serosal mapping in a porcine model

**DOI:** 10.1101/2021.08.01.454685

**Authors:** Stefan Calder, Leo K. Cheng, Christopher N. Andrews, Niranchan Paskaranandavadivel, Stephen Waite, Saeed Alighaleh, Jonathan C. Erickson, Armen Gharibans, Gregory O’Grady, Peng Du

## Abstract

Gastric disorders are increasingly prevalent, but reliable clinical tools to objectively assess gastric function are lacking. Body-surface gastric mapping (BSGM) is a non-invasive method for the detection of gastric electrophysiological biomarkers including slow wave direction, which have correlated with symptoms in patients with gastroparesis and functional dyspepsia. However, no studies have validated the relationship between gastric slow waves and body surface activation profiles. This study aimed to comprehensively evaluate the relationship between gastric slow waves and body-surface recordings. High-resolution electrode arrays were placed to simultaneously capture slow waves from the gastric serosa (32×6 electrodes at 4 mm resolution) and abdominal surface (8×8 at 20 mm inter-electrode spacing) in a porcine model. BSGM signals were extracted based on a combination of wavelet and phase information analyses. A total of 1185 individual cycles of slow waves assessed, out of which 897 (76%) were normal antegrade waves, occurring in 10/14 (71%) subjects studied. BSGM accurately detected the underlying slow wave in terms of frequency (r = 0.99, p = 0.43) as well as the direction of propagation (p = 0.41, F-measure: 0.92). In addition, the cycle-by-cycle match between BSGM and transitions of gastric slow waves in terms either or both temporal and spatial abnormalities was demonstrated. These results validate BSGM as a suitable method for non-invasively and accurately detecting gastric slow wave activation profiles from the body surface.

**Single sentence summary:** Simultaneous recordings of the stomach using serosal and body-surface electrode arrays demonstrated reliable detection of frequency and classification of propagation.

## Introduction

Disorders of gastroduodenal function encompass chronic nausea and vomiting syndromes, gastroparesis, and functional dyspepsia *(1)*. These disorders affect more than 10% of the population, imparting a significant healthcare and economic burden, and are increasing in prevalence *(2–4)*. A critical barrier to improving care remains the lack of diagnostic tests that can provide reliable and objective biomarkers of gastric dysfunction, in order to identify contributing mechanisms and personalize therapy *(5, 6)*.

Gastric motility is coordinated by an underlying electrophysiological activity called slow waves that are generated by interstitial cells of Cajal (ICC) *(7)*. In health, gastric slow waves originate at and entrain to a single pacemaker at the upper greater curvature, forming antegrade propagating wavefronts that arise approximately every 20 s and spaced 60 mm apart *(8)*. High-resolution (HR) mapping techniques have recently been employed to map slow wave activation patterns in detail at the gastric serosal surface, including in disease states *(9)*. A key finding has been that disordered gastric slow wave patterns occur frequently in patients with severe gastric symptoms, often in conjunction with ICC damage and loss, and that these abnormal patterns may occur at the nominal frequency range *(10, (11)*.

Electrogastrography (EGG) is a clinical test that assesses gastric electrophysiology by employing sparse electrode configurations non-invasively *(12)*. While EGG gas revealed consistent frequency abnormalities across several upper gastrointestinal (GI) functional disorders *(13–(15)*, it failed to achieve wide clinical adoption owing to a lack of clinical reliability, sensitivity to noise, and inadequate spatial resolution. More recently, body-surface gastric mapping (BSGM) has been proposed as novel method to detect changes in the spatial propagation of gastric slow waves non-invasively at the epigastrium *(13)*. BSGM overcomes the lack of spatial resolution that critically limited EGG, while also introducing several new spatial biomarkers in addition to more robust frequency, amplitude, and meal response profiling *(16)*. Emerging data suggests that BSGM biomarkers, and in particular abnormal slow wave direction, can offer superior symptom correlations to previous tests such as EGG and gastric emptying *(17, (18)*.

Despite the emerging potential of BSGM, there have been very few studies that demonstrated a direct correlation between gastric slow waves and EGG *(12, (19)*, and there have been no experimental studies that validated the relationship between gastric slow waves and BSGM. This validation is important because changes in the phase information at the body surface are currently being assumed to reflect true changes in the spatial organization of gastric slow waves *(17, (18)*. While a number of mathematical modeling and benchtop phantom studies have pointed to definitive and reproducible changes in body-surface potentials related to the underlying gastric slow wave activity *(20, (21)*, experimental validation remains essential before clinical translation can be accepted.

The objective of this study was therefore to definitively assess the relationship between gastric slow waves and the resultant body-surface potential profiles using simultaneous high-resolution measurements under *in vivo* conditions. We show that BSGM methods employing sufficient coverage, spatial resolution, and algorithm can be reliable in detecting gastric activation biomarkers at the body surface, including temporal profiles and changes in the direction of propagation in normal and dysrhythmic states.

## Results

### BSGM accuracy

Fourteen healthy white cross-breed weaner pigs (42 ± 2 kg) were studied. A total of 1185 cycles of gastric slow waves were assessed from 371 min (26 ±17 min per subject) of serosal data, which included approximately 227,520 individual events. Overall, 897 (76%) cycles of slow waves were characterized as normal antegrade propagation, which was observed in 10/14 (71%) of subjects we studied. Simultaneous BSGM was applied in all subjects with the serosal recordings. The flexible printed circuit (FPC) electrodes with gel-ink wells were used in eleven subjects, and the Gastric Alimetry arrays were used in three subjects.

#### Frequency match

There was a strong correlation (r = 0.99) between the frequencies measured by the serosal recordings and BSGM (**Fig. 2A**). The frequencies detected by BSGM and serosal recordings were also statistically similar across the overall, bradygastria, normal and tachygastria ranges (**Table 1)**. Bland-Altman analysis of all individual cycles of slow waves (**Fig. 2B**) demonstrated that the match between BSGM and serosal recordings was predominantly within ±0.1 cpm at the 90% confidence interval (absolute range of difference −0.3 to 0.2 cpm), across the entire range of frequencies recorded. Most activities occurred within the 3-4 cpm range (n = 753/1185) (**Fig. 2C**). When compared against the serosal recordings, the BSGM data were comparable, with BSGM non-significantly overestimating the prevalence of frequencies in the 2-3, 3-4, and >5 cpm ranges, by 41, 29, 29 cycles, respectively; while slightly underestimating the prevalence of frequencies in the <1, 1-2, and 4-5 cpm ranges, by 13, 21, 64 cycles, respectively.

**Fig 1.**
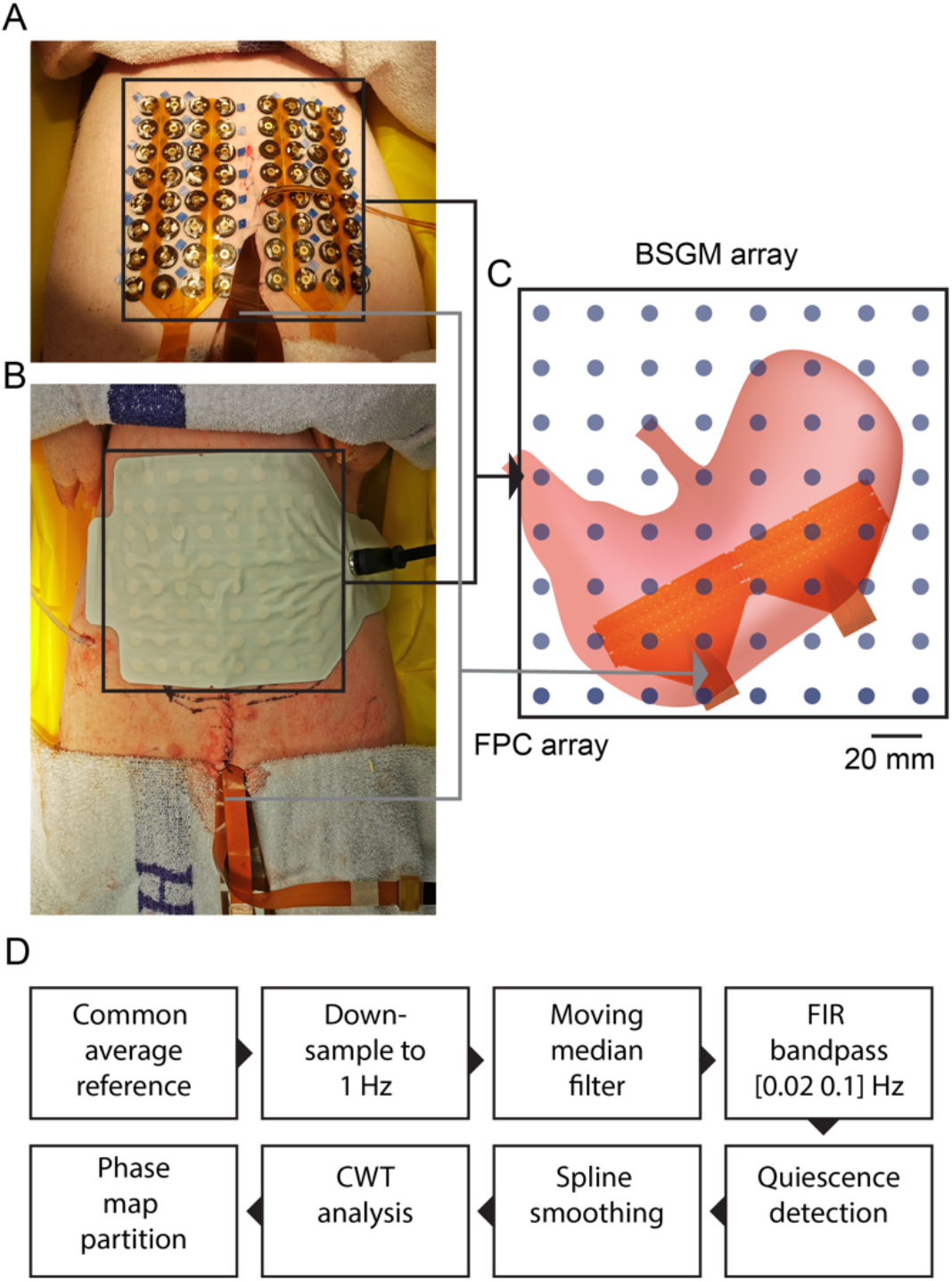
Experimental setup of simultaneous mappings from the gastric serosal surface and epigastrium. Two types of body-surface gastric mapping (BSGM) arrays were used: (A) A custom made FPC BSGM array. A rubber washer was used to hold the conductive gel between each electrode and the skin. (B) Gastric Alimetry array. Hydrogel pads formed connection between each electrode and the skin. (C) Placement of the FPC array on the gastric serosal surface. (D) Signal processing pipeline outlining the key steps in filtering, fitting procedures, transformation technique applied to BSGM signals.

**Fig. 2.**
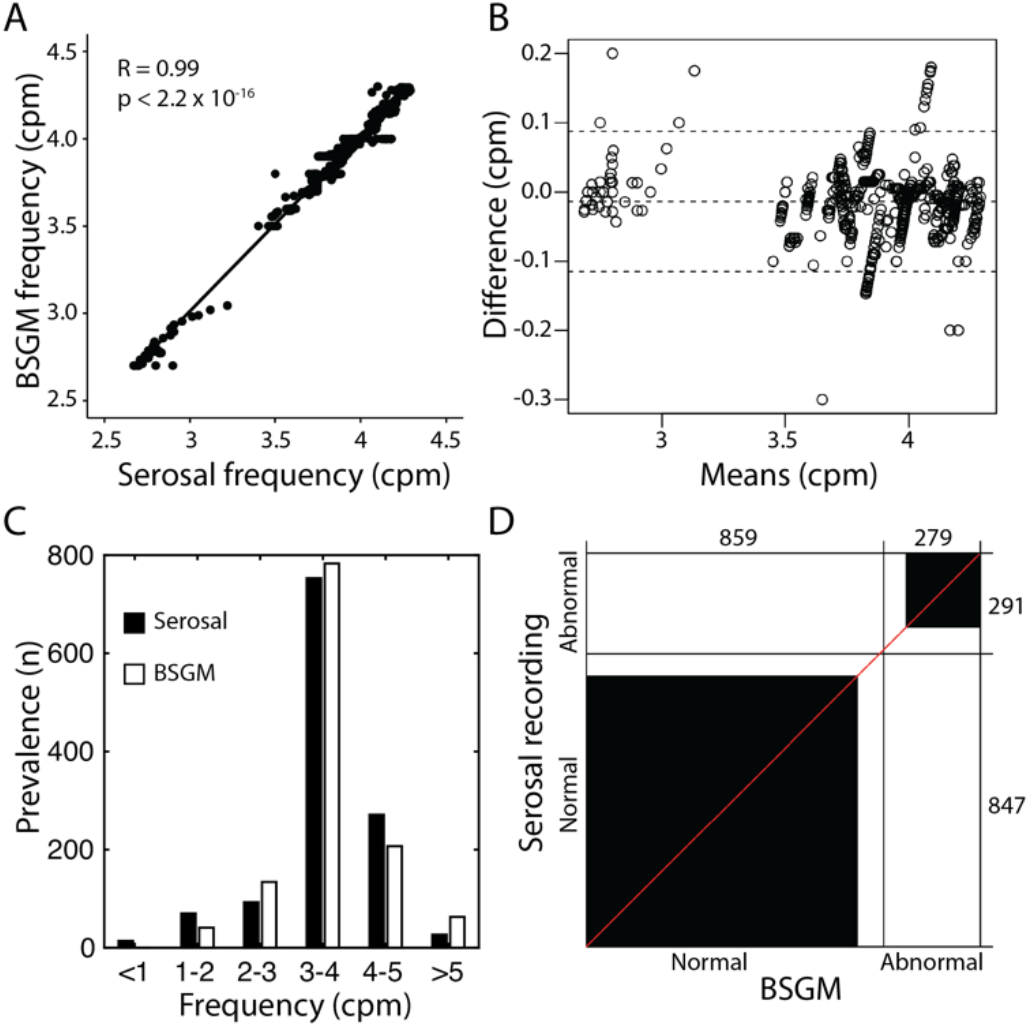
Summary statistical data of the match between serosal recordings and BSGM. (A) Correlation of the dominant frequencies, measured in cycles per minute (cpm), detected by both recordings. (B) A Bland-Altman plot illustrating the difference in the dominant frequencies. The dash lines represent the 90% confidence intervals. (C) The comparisons of the number of cycles detected by the two recordings between 1-5 cpm bands at 1 cpm intervals. (D) An agreement plot demonstrating the match between the spatial patterns detected by the BSGM when compared against the serosal recordings. Normal represents antegrade propagation and abnormal represents any significant deviations from the antegrade direction, including both initiation and conduction abnormalities.

**Table 1.**
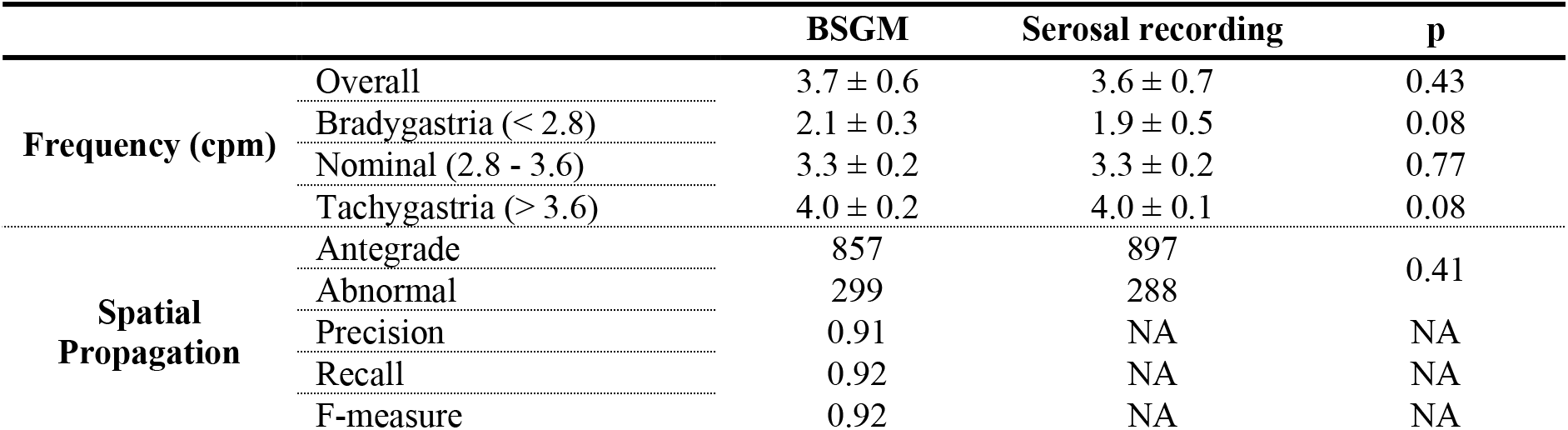
Summary statistics of the match between BSGM and serosal recordings.

BSGM also detected spatial patterns that were in good agreement with the serosal recordings (**Fig. 2D**). Out of all the activities detected by BSGM, 859 cycles were also detected as antegrade/normal and 279 cycles were detected as non-antegrade/abnormal by the serosal recordings. Conversely, out of all the activities detected by the serosal recordings, 874 cycles were also detected as normal antegrade and 291 cycles were detected as non-antegrade by the serosal recordings. The Bandiwala score was 0.81, with 1 indicating a perfect match. When the spatial propagations were analyzed independently for each type of recording, the Chi-squared analysis also demonstrated good agreement between the BSGM and serosal recordings (p = 0.41), as shown in **Table 1**. The Precision, Recall, and F-measure of the BSGM algorithm were all greater than 0.9.

#### Spatial propagation match

There was also a strong agreement between spatial patterns identified by BSGM and serosal mapping (**Fig. 2D**). Out of all slow wave cycles analyzed, 859 cycles were detected as normal antegrade by BSGM, vs 874 cycles on serosal mapping; and 279 cycles were detected as non-antegrade vs 291 by the serosal recordings (Chi-squared p = 0.41). The Precision, Recall, and F-measure of the BSGM algorithm were all > 0.9 (**Table 1**). The Bandiwala score (also the adjusted score) was 0.81, with 1.00 indicating a perfect match.

### Antegrade propagation, stable and unstable intervals

To assess the cycle-to-cycle accuracy, segments with consistent antegrade propagation longer than 300 s were analyzed, which were identified in 9/14 (64%) subjects. Antegrade propagations with a stable interval were assessed separately to those with unstable intervals, to evaluate the performance of the BSGM algorithm to variations in cycle stability. Two typical segments are presented in **Fig. 3**. In the first example (**Fig. 3A–C**), the mean interval of gastric slow waves was 15.4 ± 0.3 s over the duration of recording, which was comparable to that detected by BSGM (15.4 ± 0.9 s; p = 0.87). The difference in interval between BSGM and serosal recordings was 0.9 ± 0.6 s. Based on the electrograms and serosal activation map (**Fig. 3B,C**), all of the activities during this period were antegrade, (P1) with the BSGM phase map showing the same antegrade conduction profile throughout (**Fig. 3B,C**). In the second example (**Fig. 3D–F**), the mean interval of gastric slow waves was 20.9 ± 2.8 s over the duration of recording, with comparable values detected by BSGM (20.8 ± 2.8 s; p = 0.88). The difference in interval between BSGM and serosal recordings was 1.6 ± 1.5 s. There was a widening of intervals between 180 s and 240 s, as demonstrated by the serosal and BSGM electrograms in **Fig. 3F**. Based on the serosal activation map (**Fig. 3E**), all of the activities during this period were antegrade with the BSGM phase map again showing the same antegrade conduction profile.

**Fig. 3.**
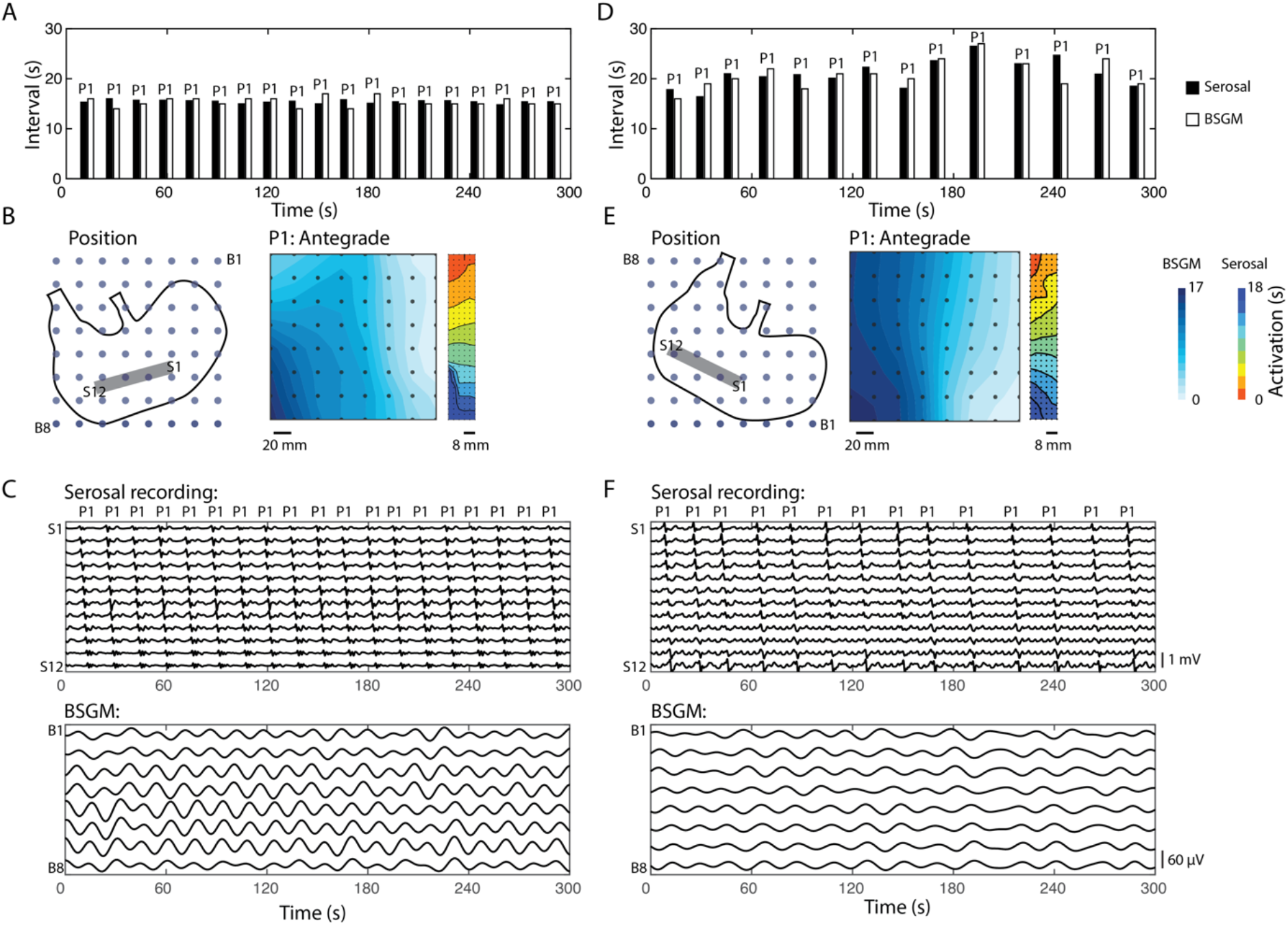
Agreement between BSGM and serosal recordings in the presence of stable antegrade propagations with stable and unstable intervals. (A) An example data showing the intervals detected by BSGM and serosal recording of a 300 s antegrade propagation with stable intervals. (B) The BSGM phase-map and serosal activation map detected during the recording in (A). (C) Sample electrograms from the serosal electrodes (S1-12) and body-surface electrodes (B1-8). (D) An example data showing the intervals detected by BSGM and serosal recording of a 300 s antegrade propagation with unstable intervals. (E) The BSGM phase-map and serosal activation map detected during the recording in (D). (F) Sample electrograms from the serosal electrodes (S1-12) and body-surface electrodes (B1-8).

### Variable propagation with nominal frequency

Persistent retrograde propagation was captured in 6/14 (43%) subjects. **Fig. 4** demonstrates the accuracy of BSGM in detecting a transition from an antegrade to retrograde spatial pattern (induced by distal gastric pacing) over a 300 s recording. In this example, except for the two cycles between 120 and 180 s, the slow wave intervals were identical during antegrade and retrograde (paced) sequences (15.9 ± 2.2 vs 16.1 ± 0.1 s; p = 0.52) (**Fig. 4A**). There was no difference in the intervals detected between two recording modalities (serosal vs BSGM; 16.9 ± 3.0 vs 16.8 ± 2.8 s; p = 0.43), with a difference of 0.7 ± 0.5 s between the two recordings. The serosal electrograms and maps (**Fig. 4B, C**) show the antegrade (P1) and retrograde (P2) sequences entraining over the entire serosal electrode array. The average velocity of the antegrade propagation was slightly faster than retrograde propagation (8.4 ± 1.2 vs 7.9 ± 1.1 mm s^−1^; p = 0.02), as indicated by the larger gaps between isochrones in the activation map of the retrograde propagation compared to the antegrade activation map (**Fig. 4B**). The change in serosal propagation pattern from antegrade to retrograde propagation was matched on a one-to-one basis by the BSGM phase maps (**Fig. 4C**). The average total phase difference of the BSGM associated with antegrade propagation was longer than the BSGM associated with retrograde propagation (14.4 ± 2.2 vs 13.2 ± 1.3 s; p = 3.8×10^−6^).

**Fig. 4.**
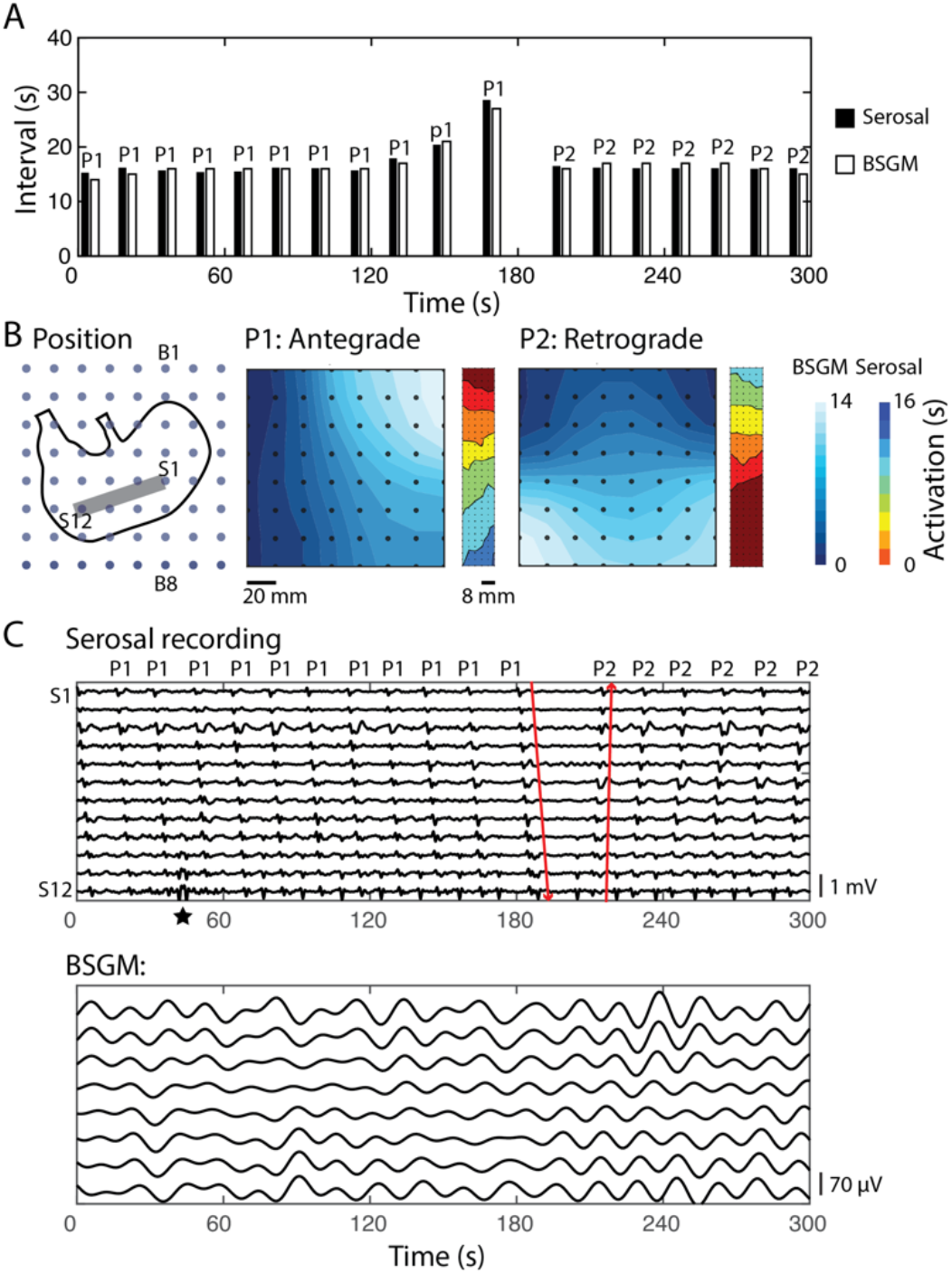
Agreement between BSGM and serosal recordings in the presence of stable frequency with a change in spatial propagation. (A) An example data showing the intervals detected by BSGM and serosal recording of a 300 s with a change in spatial patterns. (B) The BSGM phase-map and serosal activation map detected during the recording in (A). Two propagation patterns were present (P1: antegrade propagation; P2: retrograde). (C) Sample electrograms from the serosal electrodes (S1-12) and body-surface electrodes (B1-8). The star denotes the start of the stimulus that was used to invoke the retrograde propagation.

### Detection of dynamic spatiotemporal dysrhythmias

To assess the ability of BSGM to detect abnormalities during more complex dysrhythmic (dynamic) patterns, segments containing greater than 300 s of bradygastria or tachygastria based on the definition in **Table 1** with persistent abnormal propagation were identified, occurring spontaneously in 4/14 (29%) subjects. **Fig. 5** shows an example of 600 s of bradygastria with varying intervals and an antegrade propagation with a conduction block induced by a distal ectopic pacemaker. The mean interval of gastric slow waves during this period was comparable between the serosal and BSGM recordings (62.8 ± 16.8 vs 62.6 ± 18.5 s; p = 0.43), with a difference of 5.7 ± 4.6 s. The antegrade propagation occurred over a distance of 44 mm followed by an 8 mm conduction gap, which was related to a conduction block adjacent to the ectopic pacemaker arising in the middle of the mapped serosal field (**Fig. 5B**). The corresponding BSGM maps showed a comparable deviation from the normal antegrade pattern, e.g., **Fig. 3B, E**, with two early activations in the middle of the serosal electrode array (S7 in P1 of **Fig. 5C**). P1 was consistently detected until 350 s, after which the distal propagation switched to retrograde with a colliding wavefront against the proximal antegrade wavefront (P2 of **Fig. 5B**). In general, the average amplitude of the ectopic activation was lower than the colliding wavefront (0.85±1.8 vs 2.0±0.7 mV; p = 0.001), but with faster velocity (12.8±3.0 vs 11.7±2.3 mm s^−1^; p = 0.004). The average total phase difference of the BSGM associated with ectopic was shorter than the BSGM associated with colliding wavefront (11.5 ± 1.5 vs 19.7 ± 7.3 s; p= 0.006).

**Fig. 5.**
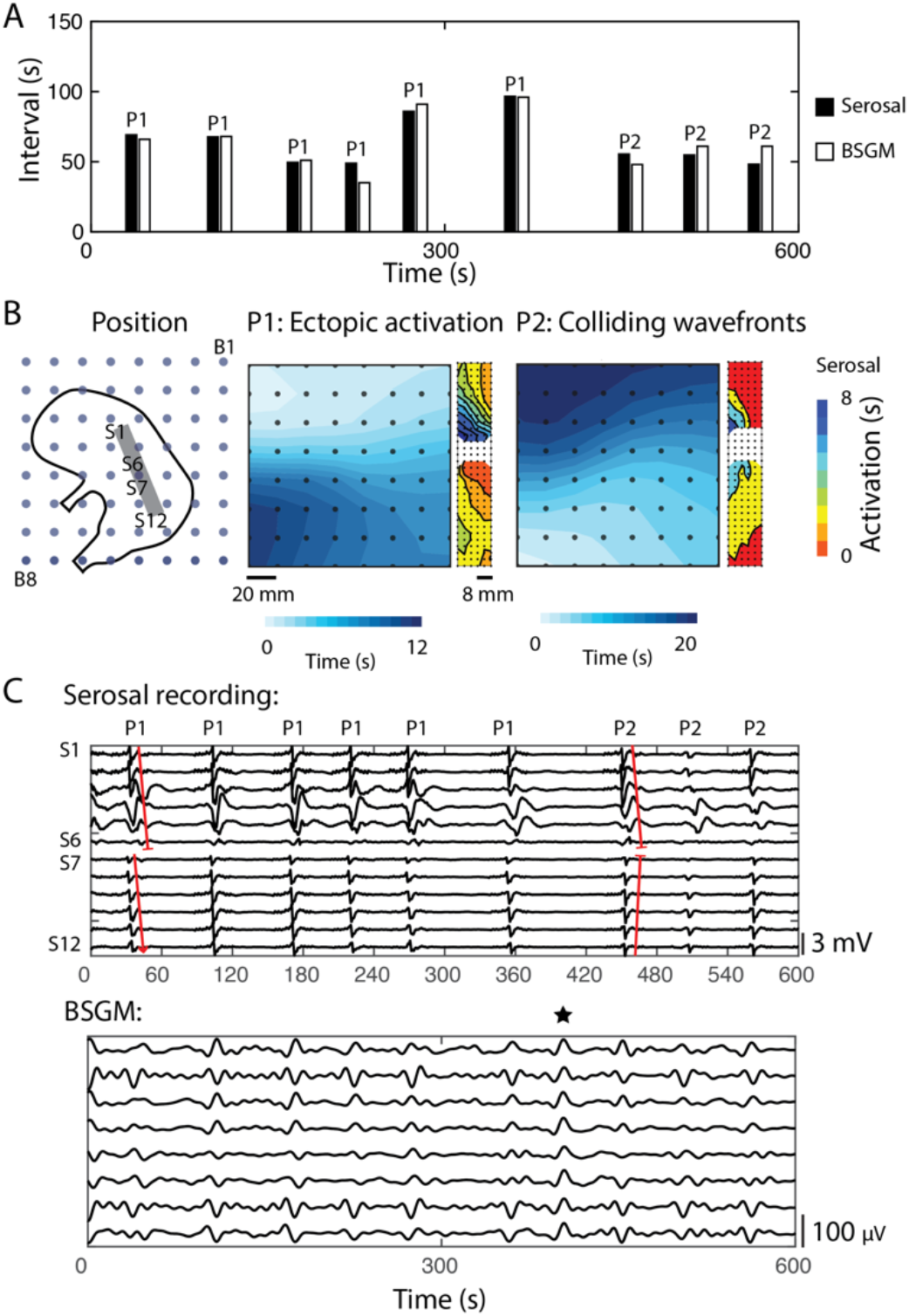
Agreement between BSGM and serosal recordings in the presence of dynamic gastric dysrhythmias exhibiting changes in both frequency and spatial propagations. (A) An example data showing the intervals detected by BSGM and serosal recording of a 600 s bradygastria with change in spatial patterns. (B) The BSGM phase-map and serosal activation map detected during the recording in (A). Two propagation patterns were present (P1: ectopic activation with a functional conduction block preventing antegrade propagation; P2: colliding wavefronts between antegrade and retrograde propagations). (C) Sample electrograms from the serosal electrodes (S1-12) and body-surface electrodes (B1-8). The star indicates an instance of noise artifact that was in the same direction of deflection in all the BSGM channels.

## Discussion

This aim of this study was to definitively evaluate the relationship between gastric slow waves and body-surface potentials. The data provide clear evidence that BSGM, when performed with the specific algorithms, is a suitable method for non-invasively measuring the frequency and direction of propagation of gastric slow waves. The data demonstrated that the phase maps produced by BSGM from recordings at the abdominal surface were in consistently strong agreement with the frequency and pattern of slow wave propagation in the stomach, including during inconsistent variations in wave intervals and propagation directions. Together, these data validate BSGM as a non-invasive tool for reliably detecting gastric slow wave biomarkers.

This data is important because BSGM is proposed as an emerging tool for clinically evaluating gastric function *(13)*. Early studies have shown promising correlations between upper GI symptoms and the direction of propagation measured from the body surface, including in functional dyspepsia, gastroparesis, and nausea and vomiting in childhood *(18)*, which were not apparent for frequency metrics derived from conventional EGG. Further studies are needed to validate these early correlations, which can now be enabled with BSGM.

The interpretation of BSGM warrants a more detailed understanding of the mechanisms of transmission of slow waves from the stomach to the body-surface. Based on biophysical principles, a dipole presenting the net summation of gastric slow waves may be used to calculate the resultant body-surface potential *(22)*. However, such an assumption is only valid when the wavefronts are generally propagating in the same direction *(20)*. Whether the waves are consistently antegrade or retrograde, the direction of the dipole would generally be in agreement with the slow wave directionality, which was evident in **Figs. 3** and **4**. However, as wavefronts become disorganized or fractionated, there would be less agreement between the direction of the BSGM and the underlying gastric slow waves, as shown in **Fig. 5**. Therefore, instability of phase-maps can be indicative of dynamic underlying propagation patterns. Furthermore, BSGM has not yet been validated for detection of localized changes in the speed of slow waves. Time differences between the BSGM phase maps and serosal maps were evident, which was likely due to the change in direction rather than increase or reduction of speed of slow wave propagation, with projection onto the body surface.

Some caution must be noted when extrapolating the results to clinical use. First, the animal subjects were under general anesthesia in a supine position and immobile. The respiration was controlled at a known rate using an external ventilator, which enabled straightforward elimination of respiratory noise from the signals. The immobility of the subjects also meant the signals were not contaminated by any other movement noises. In practice, artifacts due to movements and respiratory will need to be accounted for by an accelerometer and filtered using a filtering scheme such as the linear minimum mean square error (LMMSE) method *(23)*. Second, in this animal study, positioning of the body surface array directly over the stomach was readily achieved by visual reference at the time of laparotomy. In clinical practice, other methods of reliably locating the array over the stomach will be required to ensure optimal signal quality *(24)*. Human gastric slow waves also occur with a closer wave separation than in pigs due to the slower conduction velocity in the corpus and proximal antrum (~6 cm vs ~14 cm wave separations) *(25)*, which may necessitate further tuning of current algorithms for clinical applications *(20)*.

In addition, while the weaner pig model presents a useful range of dysrhythmias that are broadly comparable to common human dysrhythmic patterns *(26)*, abnormalities in patients with severe gastric dysfunction may be more complex and unstable than those represented here, or are sometimes constrained to a localized gastric regions *(27)*. The spatial propagations in the present study were broadly categorized as either normal or abnormal, but further refinement of the accuracy of BSGM of detecting the different subcategories of gastric dysrhythmias, e.g., initiation and conduction dysrhythmias, is likely to continue in future *(20)*. Ultimately, however, resolutions achievable at the epigastrium will never be as granular as those achieved at the serosal surface, due to the volume conduction and summation of body surface potentials *(21)*. Nevertheless, it was clearly here shown that robust classification of overall slow direction was achievable, and moreover that it is possible to detect some more complex patterns such as competing pacemakers or colliding waves. Gastric dysrhythmias in patients should therefore also be robustly detectable at the body surface when there is a deviation from antegrade propagation, while potentially also showing corresponding frequency instabilities on spectral analytics *(13, 18, 28)*.

A pertinent question is whether the changes in gastric slow waves, as detected by BSGM in this study, would necessarily relate to changes in gastric motility. The current study was not designed to address this question, but several animal studies have previously shown that when retrograde propagation occurs at frequencies close to the normal range, motility follows *(29, 30)*. Although this connection has not yet been made in humans, emerging correlations between symptoms and retrograde propagation indicate a plausible pathophysiological connection *(31)*.

Further development of BSGM will focus on enhancing its value as a clinical tool. Scalable and clinically intuitive devices are need that can offer the ability to monitor gastric electrophysiology and meal responses continuously over several hours. Such systems could provide a more comprehensive evaluation of gastric function than transit time (i.e. gastric emptying testing), which has correlated only weakly with symptoms, and has recently been criticized as labile *(6, 32)*. Furthermore, BSGM can be combined with different types of provocative tests, such as test meals and water load, as well as interventional measures such as pharmaceuticals and neuromodulation *(33, 34)*, to improve sensitivity and clinical efficacy. Real-time symptom recording could also be performed during the recording to further establish the link between changes in gastric slow waves, subject condition and symptom scores. Given the large number of patterns generated by BSGM, machine learning may also help to streamline the analysis procedure in future, by grouping together similar patterns and identifying features that are relevant to different symptoms and/or phenotypes of diseases *(35, 36)*. Finally, relating the body-surface potentials in the inverse manner will help to potentially pinpointing locations of persistent ectopic pacemakers to potentially guide electroceutical and ablation therapies *(37, 38)*.

In conclusion, this study definitively validates BSMG as a reliable diagnostic strategy to non-invasively detect changes in gastric slow waves. Translation can now proceed with confidence to define the clinical potential of the novel biomarkers provided by BSGM, including their relationship to symptoms and ability to impact clinical decisions.

## Material and Methods

Ethical approval was granted by the University of Auckland Animal Ethics Committee and the guide for the care and use of laboratory animals was followed *(39)*. As a large monogastric mammal species, white cross-breed weaner pigs are an appropriate translational model for studying gastrointestinal electrophysiology and offers a close approximation to human physiology *(25)*. An incidence of dysrhythmias of 15% in the weaner pig model, based on a previously published mapping study, was used in a power calculation to determine the number of subjects required *(26)*, with an α error probability of 0.05 and a power of 0.80, which resulted in a sample size of 14 subjects and a total of 386 cycles of gastric slow waves. Each subject was induced with Zoletil (tiletamine HCl 50 mg mL-1 / zolazepam HCl 50 mg mL^−1^) and maintained via isoflurane (2.5 – 5%) with an oxygen flow of 400 mL within a closed-circuit anesthetic system. Vital signs were monitored continuously via a femoral arterial line, rectal thermometer, and capnography. Gastric pacing was also available to augment the rate of retrograde gastric activation sequences in the experimental model to meet this target, if required *(40)*.

### Recording Protocol

A summary of the recording setup is illustrated in **Fig. 1A**. The abdominal skin of the pig was shaved with surgical clippers (CareFusionVernon Hills, IL, USA) and prepared using Nuprep (Weaver and Company, Aurora, CO, USA). A midline laparotomy of approximately 100 mm in length was performed to access the stomach. A flexible printed circuit (FPC) electrode array (192 channels at 4 mm inter-electrode spacing) (FlexiMap gen3 v5, New Zealand) was gently positioned onto the corpus of the stomach to record gastric slow waves *(41)*. Each internal FPC array was connected to an ActiveTwo acquisition system (BioSemi, Netherlands) via a 32-way ribbon cable. The ActiveTwo system was then connected to a recording Laptop with a fiber-optic to USB connection. A custom recording software (written in LabView v8.2, National Instruments, TX, USA) was used to record the serosal gastric slow waves at a sampling frequency of 512 Hz. In some studies, a gastric stimulation (pacing) device was connected to a pair of pacing electrodes placed distal to the FPC arrays on the stomach *(40)*. The incision was then sutured closed for the duration of the experiment. Where required, a stimulation protocol (amplitude: 4 mA; pulse-width: 200 ms; interval: 16 s) was administered to induce retrograde propagation in the stomach.

A custom BSGM FPC array with 8×8 electrodes was developed. It was designed in order to integrate with a disposable ECG dot (Kendall™/Covidien Medi-Trace® 200 Series, Medi-trace Inc., USA) through a 4 mm diameter hole using a tinned snap-connector, as shown in **Fig. 1B**. In three subjects, a comparable custom conformable BSGM array (8×8 Ag/AgCl electrodes) printed on thermoplastic substrate prepared with conductive hydrogel discs and an adhesive laminate (Alimetry, New Zealand), was employed. In both cases, the end of the array was connected using two 32-way ribbon cables to the BioSemi system in the same way as the serosal FPC array.

### Analysis Protocol

The serosal/reference data was analyzed in the Gastric Electrical Mapping Suite (GEMS v1.7, The University of Auckland, New Zealand) via MATLAB (R2018b, Mathworks, MA, USA)*(42)*. The activation time of each gastric slow wave was automatically marked then grouped into discrete wavefronts (cycles), followed by manual review and correction by at least two investigators. Normal antegrade propagation was classified as propagation that followed the organo-axis of the stomach from the fundus towards the antrum *(25)*, with directional deviations from antegrade propagation defined as abnormal *(43)*. The average amplitudes, velocity and time interval from the preceding cycle were quantified for every cycle of analyzed slow waves.

The body-surface data was analyzed using a proprietary algorithm coded in Python (version 3.7) (Alimetry, New Zealand), which consisted of a series of processing and spatiotemporal analysis modules illustrated in **Fig. 1D**. The common average reference (CAR) was subtracted from all BSGM channels, and the signals were then iteratively down-sampled from 512 to 1 Hz. The down-sampled signals were filtered first by applying a central moving median filter window size of two minutes] *(44)*, then a FIR bandpass filter [0.02-0.1], and finally smoothed with a polyharmonic spline method *(45)*. The continuous wavelet transforms (CWT) using a morlet wavelet was used to analyze the spectral information *(46)*, followed by a moving mean filter to remove any remaining noise. Quiescence, or periods of no slow wave activity, was determined by implementing an adaptive threshold on the spectral decomposition of each channel, periods below the threshold were eliminated form further analysis.

The CWT transform was applied to calculate the phase of individual BSGM channels. Using the phase information, the data was first separated into wave intervals and allocated over the the 8 × 8 BSGM array *(47)*. An iterative matching scheme using a Pearson correlation coefficient with a threshold of 0.6 was implemented to group similar spatial propagations together.

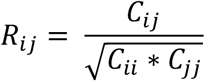

where R denotes the correlation coefficient matrix and C denotes the covariance matrix for the signal electrode located at i,j in the BSGM array. Every wave was assigned a group unless the threshold was not reached, in which case a new group was created.

Frequency and spatial propagation of each cycle of body-surface was reported in the same manner as the serosal data. Normal and abnormal propagations were defined based on the relative position and orientation of the BSGM array to the stomach, which was labelled on the abdominal skin prior to closing the incision. To quantify the performance of spatial characterizations, we defined the *true positive* (TP) as when the same pattern was detected by both the serosal and BSGM recordings, *false positive* (FP) as when different patterns were detected by the serosal and BSGM recordings, *false negative* (FN) as when the BSGM missed a pattern detected by the serosal recordings. The following metrics were used to quantify the accuracy of the BSGM analysis:

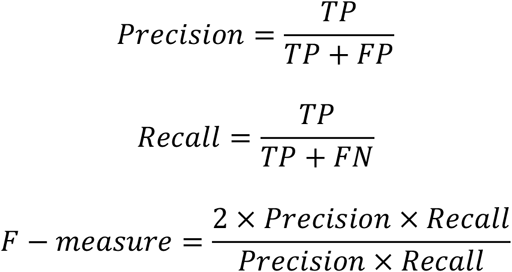

### Statistical Analysis

A Student’s t-test was performed to compare the difference between the frequency measured from the reference data and body-surface data. The null hypothesis was that there was no difference between the two datasets. The Bandiwala score was calculated to measure the agreement of the spatial patterns between the two recordings *(48)*. A Pearson’s Chi-squared test was performed a measure of goodness of fit for the normal and abnormal categories. The null hypothesis was that the frequency distribution of the body-surface characterized patterns were consistent with the distribution of the reference data characterized patterns. A p of < 0.05 was deemed to show statistical significance between the two datasets.

## Acknowledgements

We thank Linley Nisbet for her assistance with the animal studies. This work and authors were funded by a Rutherford Discovery Fellowship from the Rutherford Foundation Trust and Prime Minister’s Emerging Scientist Prize administered by the Royal Society Te Apārangi, as well as a feasibility grant from the MedTech CoRE.

